# Cancer Cell Lewis X Plays a Minor Role in NK Cell Immune Evasion

**DOI:** 10.64898/2026.08.25.746752

**Authors:** Remi Hatinguais, Alba Gabarroca Garcia, Amouretty Agard, Victor Lorrain, Priscilla D.A.M. Heijnen, Sandra J. van Vliet

**Affiliations:** Department of Molecular Cell Biology and Immunology, Amsterdam UMC, Vrije Universiteit Amsterdam, Amsterdam, the Netherlands; Cancer Biology and Immunology, Cancer Center Amsterdam, Amsterdam, the Netherlands; Cancer Immunology, Amsterdam Institute for Immunology and Infectious Diseases, Amsterdam, the Netherlands

## Abstract

Production of aberrant glycans by cancer cells constitutes a key immunosuppressive strategy to avoid destruction by immune cells. Although sialic acid-containing glycans are known to dampen the activation of lymphocytes, including Natural Killer (NK) cells, the role of fucose-containing glycans remains poorly characterized. In this work, we explored the role of Lewis X (LeX) in cancer cell-NK cell interactions. We induced ectopic expression of *FUT9*, an α1-3/4-fucosyltransferase, in two colorectal cancer cell lines and showed this enzyme only synthesized LeX structures but not sialyl-LeX. *FUT9* introduction was not associated with altered MHC class I surface expression, nor with CD2 (which has been proposed as a receptor for LeX) binding to cancer cells. By inhibiting fucosylation we could demonstrate that CD2 binding was furthermore independent of surface fucosylated glycans in three independent cell lines. Lastly, FUT9/LeX had a limited role in cancer cell destruction and expression of activation markers by NK cells. Overall, our study suggests that, unlike sialylated glycans, α1-3/4-fucosylated glycans have limited impact on cancer cell evasion of NK cell-mediated destruction.

## Introduction

Alterations in glycosylation are a universal hallmark of cancer cells. These aberrant glycans are key players in pathology as they alter cancer cell properties, notably by increasing proliferation and resistance to treatment, as well as promoting immune evasion (Hatinguais *et al*, 2026). Several mechanisms of glycan-mediated immunomodulation of anticancer immune responses exist. Glycosylation directly affects the stability and function of immune-related proteins, such as checkpoint inhibitors and cytokine receptors (Hatinguais *et al*., 2026). In addition, tumor-associated glycans can trigger inhibitory receptors expressed by immune cells (Hatinguais *et al*., 2026; van Vliet & van Kooyk, 2025). For example, sialic acid-containing glycans engage receptors of the Siglec family, which exert direct immunosuppressive effects on myeloid and lymphoid cells (Hatinguais *et al*., 2026; Hudak *et al*, 2014; van Vliet & van Kooyk, 2025). Although a large body of work has focused on characterizing the role of the sialic acid-Siglec axis in immune evasion by tumor cells (Hudak *et al*., 2014; van Vliet & van Kooyk, 2025), the role of other glycan classes, including fucose-containing glycans, remains poorly understood.

Fucosylation, the addition of fucose moieties to glycoproteins and lipids is performed by 13 fucosyltransferases in human, each with a characteristic linkage specificity (Blanas *et al*, 2018; Liner & Laubli, 2026). Amongst the different glycans they produce, FUT1-7, and FUT9-11 synthesize Lewis antigens, *i*.*e*. terminal fucosylation with an α1-2, α1-3, or α1-4 linkage (Bitaraf *et al*, 2026; Blanas *et al*., 2018). FUTs and the Lewis antigens they produce are of particular relevance in cancer as they are able to promote metastasis, increase resistance to chemotherapy, or reprogram cancer cells toward a stem cell-like phenotype (Bitaraf *et al*., 2026; Blanas *et al*., 2018; Liner & Laubli, 2026). In some cases, Lewis antigens expressed by cancer cells can also improve the anticancer immune response. For instance, increased FUT3- or FUT6-mediated fucosylation of Death Receptor 5 (DR5) sensitizes cancer cells to lysis by tumor necrosis factor (TNF)-related apoptosis-inducing ligand (TRAIL) (Zhang *et al*, 2019) .

Lewis antigens can also be recognized by innate immune receptors. For instance, Lewis antigens X and Y (LeX, LeY) are ligands for the C-type lectin receptor DC-SIGN (Meyer *et al*, 2005; van Die *et al*, 2003), which initiates an anti-inflammatory response in the context of parasitic infections (Gringhuis *et al*, 2014). Interestingly, LeX (also known as CD15) has been proposed as a ligand for CD2, an activating receptor in T cells and Natural Killer (NK) cells (Sabry *et al*, 2011). Mechanistically, *FUT4* expression increased cancer cell susceptibility to NK cell cytotoxicity, which could be reverted by a blocking anti-CD2 or anti-CD15 antibody (Sabry *et al*., 2011). Therefore, LeX appears to promote NK activation, potentially in a CD2-dependent manner.

Overall, more research is needed to decipher the role of Lewis antigens in immune evasion by cancer cells. In this study, we aimed to further characterize the involvement of LeX in the interaction of cancer cells with NK cells. Namely, we investigated how LeX affected CD2 and NK activity. In contrast to a prior study (Sabry *et al*., 2011), our results show that CD2 binding to cancer cells appears unrelated to surface Lewis antigens, and that LeX does not affect cancer cell susceptibility to NK-mediated killing.

## Material and methods

### Cell line origin and maintenance

Colo320, Colo205, and LS174T were a kind gift from R. Fijneman (Netherlands Cancer Institute, Amsterdam) (26172302). Hutu80 cells were a kind gift from J.P. Medema (Amsterdam UMC). All cell lines were maintained in RPMI (Gibco) supplemented with 5% Fetal Calf Serum (FCS, Biowest), 1000 u/mL Pencillin/Streptomycin, 2 mM L-Glutamine, and 25 mM HEPES (all Gibco). Cells were routinely tested for mycoplasma contamination.

### Construction of Colo320 and Hutu80 glycovariants

The pRP-hPGK-V5-FUT9 plasmid was ordered from VectorBuilder (Chicago, USA) (Figure S1). Prior to cell transfection, plasmid was linearized with ScaI-HF (NEB), and gel purified with GeneJET kit (ThermoScientific). Mock cell lines were transfected with gel-purified pRP-hPGK plasmid from which *FUT9* had been excised using ClaI and AflII (both NEB), and gel purified (Fig. S1, and not shown). For transfection, 100,000 wild-type Colo320 and Hutu80 cells were plated in a 6-well plate and transfected with linearized plasmid using Lipofectamine LTX Plus (Invitrogen) according to the manufacturer’s protocol. 24 h after transfection, cells were selected with G418 selection (1000 μg/mL for Colo320 and 800 μg/mL for Hutu80). LeX-positive cells stained as described below were sorted using the FACS Aria (BD). LeX expression was regularly checked throughout the study, and, when necessary, CD15^+^ enrichment was performed using CD15 Dynabeads (ThermoScientific).

### Lectin profiling, Lewis antigen and CD2-Fc staining

All steps were performed in HBSS supplemented with 0.5% Bovine Serum Albumin (BSA, Roche). 50,000-100,000 cells were seeded into a 96-well V-bottom plate and stained with LiveDead Far Red Fixable Viability Dye (1:5,000, Invitrogen) and incubated for 20 min on ice in the dark. Cells were washed and subsequently incubated with biotinylated lectins, anti-Lewis antigens antibodies, or CD2-Fc (Table 1) for 30 min on ice in the dark. After washing, cells were stained with streptavidin-PE (1:400, Jackson Immunoresearch), streptavidin-APC (1:400, Jackson Immunoresearch) anti-mIgM-APC (1:200, Jackson Immunoresearch) or anti-hIgG Fc-AF647 (1:400, Biolegend) for 30 min on ice and then washed. Cells were fixed using 1% formaldehyde for 20 min at room temperature in the dark and analyzed by flow cytometry on an Attune NxT. The data were analyzed using FlowJo v.10.10.

**Table 1.** – List of plant lectins and antibodies used in this study (see supplementary information for abbreviation list)

| Lectin/Antibody | Specificity/Clone | Source | Working concentration |
| --- | --- | --- | --- |
| ConA | Diantennary *N*-glycans, high mannose | Vector Labs | 5 µg/mL |
| HPA | Tn antigen | Vector Labs | 5 µg/mL |
| LEA | LacNAc | Vector Labs | 0.5 µg/mL |
| LTA | α1-3/4 fucose | Vector Labs | 5 µg/mL |
| MAL-I | α2-3 Sia-Gal-GalNAc (*N*-glycans) | Vector Labs | 5 µg/mL |
| MAL-II | α2-3 Sia-Gal-GalNAc (*O*-glycans) | Vector Labs | 5 µg/mL |
| PHA-L | Tetra-antennary (*N*-glycans) | Vector Labs | 2 µg/mL |
| PNA | T antigen/Core 1 (*O*-glycans) | Vector Labs | 5 µg/mL |
| SBA | α-/β-GalNAc | Vector Labs | 5 µg/mL |
| SNA | α2-6 Sialic acid (*N*-glycans) | Vector Labs | 5 µg/mL |
| UEA-I | α1-2 fucose | Vector Labs | 5 µg/mL |
| AAL | α1-3/4 and α1-6 fucose | Vector Labs | 5 µg/mL |
| Anti-LeX | MC480 | BD Biosciences | 1 µg/mL |
| Anti-sialyl-LeX | SLEX | BD Biosciences | 2.5 µg/mL |
| CD2-Fc | n/a | R&D Systems | 5 µg/mL |

### Glycosylation inhibitor treatment

200,000 cells were seeded into a 6-well plate and allowed to adhere for at least 2h prior to treatment with the glycosylation inhibitors 2-fluoro-peracetyl-fucose (2F-PF, 200 μg/mL, MedChemExpress), Kifunensine (10 μg/mL, R&D Systems), BenzylGalNAc (500 μM, MedChemExpress) or Gen123346 (500 μM, Abcam). Dimethylsulfoxide (DMSO)-treated cells were used as control. After 3 days of treatment, cells were harvested and stained with lectins, anti-Lewis antibodies or CD2-Fc, as described above.

### IFN-γ and MHC class I staining

100,000 Colo320 or Hutu80 glycovariants were seeded into a 24-well plate and incubated overnight at 37°C 5% CO2. The next day, medium was replaced and cells were treated with 0.5 ng/mL rhIFN-γ (Immunotools) for 24 h, after which cells were harvested. Viability staining was performed as described above, MHC class I was stained using anti-HLA Class I-AF700 (1:100, clone W6/32, Biolegend) for 30 min. Cells were fixed as described above, acquired on the Attune NxT and data were analyzed using FlowJo v.10.10.

### NK cell isolation

Peripheral blood mononuclear cells (PBMCs) from healthy donors were obtained from buffy coats (Sanquin Amsterdam, The Netherlands) by Ficoll gradient centrifugation. NK cells were isolated using the MojoSort Natural Killer Negative selection kit (Biolegend) according to the manufacturer’s protocol. After isolation, NK cells were resuspended at 5×10^6^ cells/mL in RPMI 10% FCS supplemented with 50 IU/mL rhIL-2 (Immunotools) and incubated overnight at 37°C 5% CO2 . The next day, NK cells were harvested and counted prior to assays.

### NK cytotoxicity assay

30,000 cancer cells were seeded into a 96-well plate and incubated overnight at 37°C 5% CO2. The following day, the medium was removed and 30,000 NK cells were added in 200 μL RPMI 10% FCS. After 4 h or 24 h, CellTiterBlue (CTB, Promega) metabolization was used as a proxy for cell viability. Cancer cells without NK cells were used as a 100% viability controls, and NK cells only were included as a control for NK-mediated CTB metabolization. Fluorescence (590 nm, F590) was measured on a Synergy HT (BioTek) after 2-4 h at 37°C 5% CO2. Percentage of killing was calculated by the formula % *killing* = 100 − 100 * 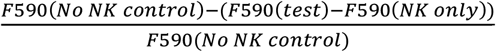. All conditions were performed in technical duplicates or triplicates and with 5-8 independent healthy donors.

#### NK activation marker assay and staining

100,000 cancer cells were seeded in a 24-well plate and incubated overnight at 37°C 5% CO2. The next day, the medium was discarded and 100,000 NK cells were added in RPMI 10% FCS containing anti-CD107a-BV421 (1:100, Biolegend). 4 h before the end of the assay, GolgiPlug (1:1000, BD Biosciences) and GolgiStop (1:1500, BD Biosciences) were added to the wells.

After a total of 4 hours or 24 hours, cells in suspension and adherent cell were detached with TrypLE (Gibco), harvested and pooled. Cells were resuspended in PBS 0.5% BSA, and incubated with LiveDead Far Red Fixable Viability Dye (1:5,000, Invitrogen) and Human TruStain FcX Fc-block (1:200, Biolegend) for 20 min on ice in the dark. Cells were washed and then stained with an antibody cocktail (Table 2) and incubated on ice for 30 min. Cells were washed, fixed with 1% formaldehyde for 20 min at room temperature in the dark. Cells were washed and subsequently permeabilized with PBS 0.5% BSA 0.3% saponin 2 mM EDTA and Human TruStain FcX Fc-block (1:200, Biolegend) for 20 min on ice in the dark. After washing, cells were stained intracellularly (Table 2) for 30 min at room temperature in the dark. Cells were washed twice and measured by flow cytometry on a Northern Light 3 laser instrument.

**Table 2.**
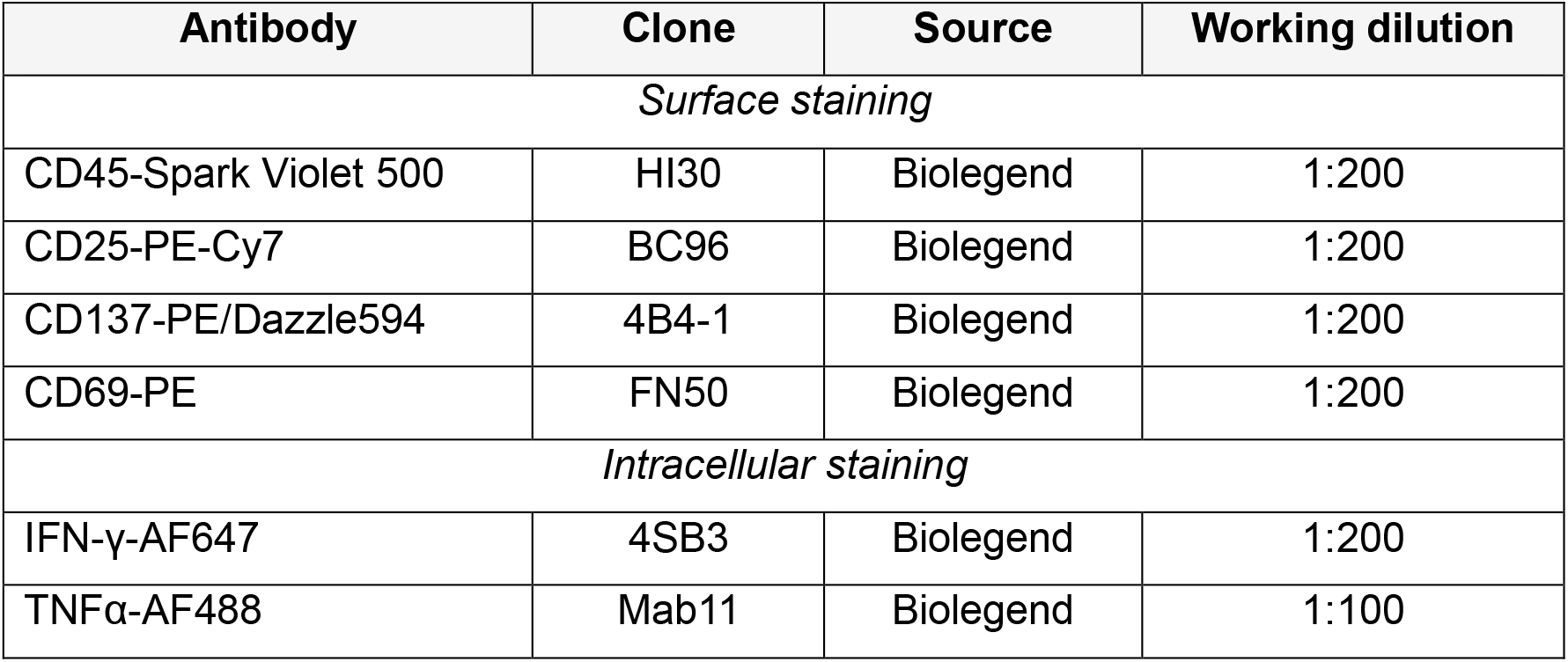
– Antibodies used to assess NK activation markers.

| Antibody | Clone | Source | Working dilution |
| --- | --- | --- | --- |
| *Surface staining* |  |  |  |
| CD45-Spark Violet 500 | HI30 | Biolegend | 1:200 |
| CD25-PE-Cy7 | BC96 | Biolegend | 1:200 |
| CD137-PE/Dazzle594 | 4B4-1 | Biolegend | 1:200 |
| CD69-PE | FN50 | Biolegend | 1:200 |
| *Intracellular staining* |  |  |  |
| IFN-γ-AF647 | 4SB3 | Biolegend | 1:200 |
| TNFα-AF488 | Mab11 | Biolegend | 1:100 |

### Statistics

Statistics, as indicated in the figure legends, were calculated using GraphPad Prism v.10.6.0.

## Results

### Characterization of FUT9-expressing human CRC cells

In order to further explore the role of LeX, we selected to focus on FUT9, as this fucosyltransferase selectively synthesizes LeX and not sLeX (Mondal *et al*, 2018). We transfected two colorectal cancer cell lines, Colo320 and Hutu80, both belonging to the aggressive subtype consensus molecular subtype 4 (CMS4) (Guinney *et al*, 2015; Linnekamp *et al*, 2018) with a *FUT9-*encoding plasmid (Fig. S1). We decided to express *FUT9* under the control of the human phosphoglycerate kinase (PGK) promoter (Fig. S1), which is expected to induce a constitutive low-to-moderate gene expression, resulting in more physiologically-relevant levels than viral promoters (Qin *et al*, 2010). After selection, we observed that both Colo320-FUT9 and Hutu80-FUT9 expressed high levels of surface LeX but not sialyl-LeX (sLeX) (Fig. 1A), as expected (Mondal *et al*., 2018). Neither LeX nor sLeX could be detected at the surface of Mock-transfected cells (Fig. 1A), confirming that all LeX detected was the product of FUT9.

**Figure 1.**
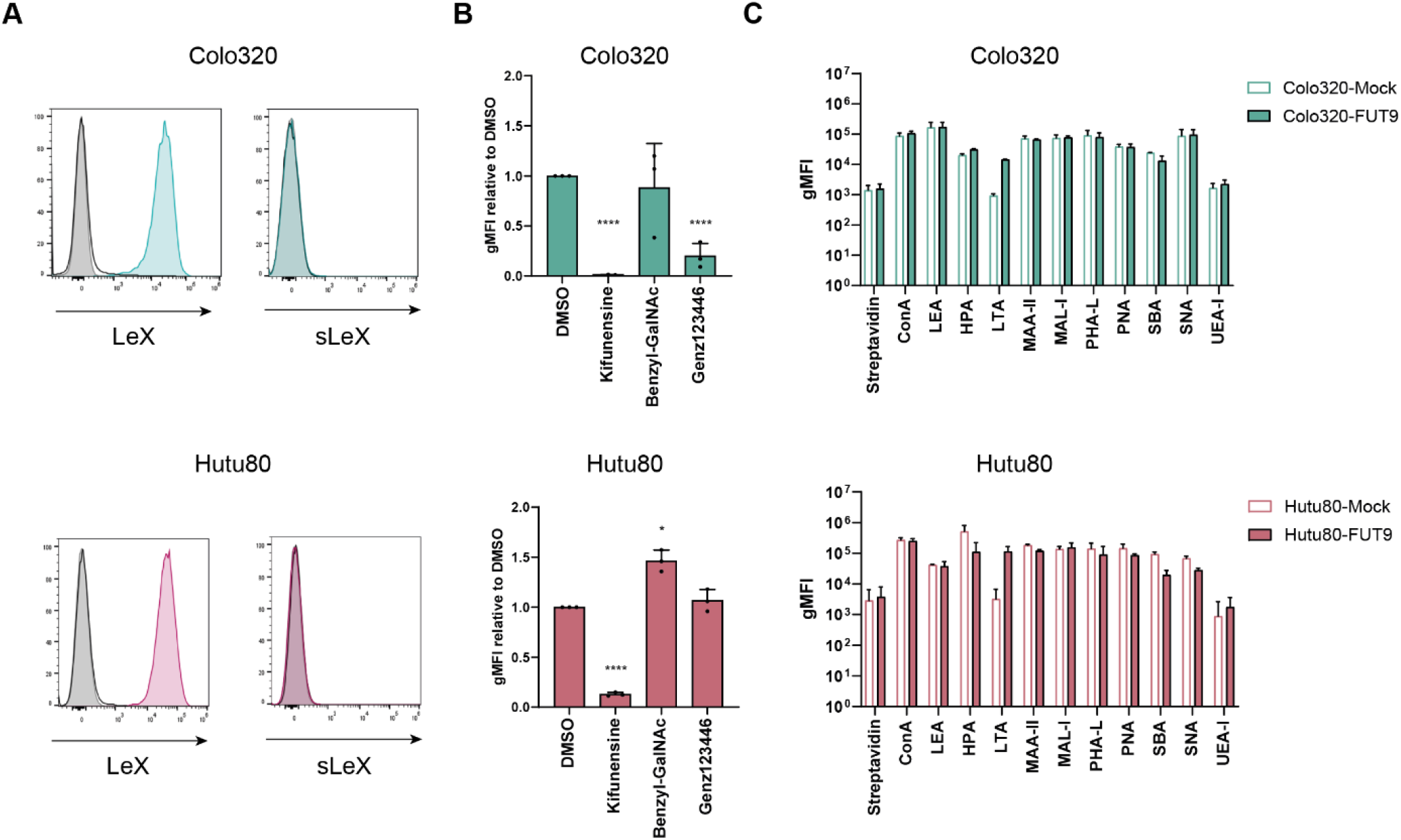
– Glycosylation characterization of FUT9-expressing cells. **A**, Colo320 (top, green) or Hutu80 (bottom, red) glycovariants were stained for LeX or sLeX and analyzed by flow cytometry. One representative experiment is depicted. Grey, streptavidin-stained cells; Black, Mock-transfected, Filled, FUT9*-*expressing cells. **B**, Colo320-FUT9 and Hutu80-FUT9 were incubated 72h with glycosylation inhibitors, and then stained for LeX and analyzed by flow cytometry. LeX surface staining is expressed as a ratio of geometric mean fluorescence intensity (gMFI) compared to DMSO-treated control. N = 3. Normality was verified by Shapiro-Wilk and statistical differences were calculated by one sample t test. *, p < 0.05; ***, p < 0.001. **C**, Surface glycans of Colo320 and Hutu80 glycovariants were stained with a panel of lectins and measured by flow cytometry. N = 3 Statistical differences were calculated by Multiple Mann-Whitney adjusted for FDR.

FUT9 can fucosylate *O-*glycans, *N-*glycans, and glycolipids (Mondal *et al*., 2018) and we set to determine on which type of glycans LeX was present in our model. In both cell lines, treatment with mannosidase inhibitor kifunensine significantly reduced surface expression of LeX (Fig. 1B), indicating that most LeX epitopes are carried by *N-*glycans. In Colo320, we also observed a decrease in surface LeX after treatment with the glycolipid inhibitor Genz133456, suggesting that FUT9 also fucosylates glycolipids in this cell line (Fig. 1B). In contrast, no reduction in LeX staining was observed after treatment with Benzyl-GalNAc (Fig. 1B), showing that FUT9 does not produce LeX-carrying *O*-glycans in Colo320 or Hutu80.

Because of the ectopic expression of *FUT9* in our model, and the possibility of competition for glycosylation sites between glycosyltransferases, we assessed the overall surface glycan landscape in our glycovariants using a panel of plant lectins (Fig. 1C). In accordance with our initial results (Fig. 1A), we found that Mock-transfected cells did not express surface α1-3/4-fucose, as indicated by the absence of staining by the LTA lectin (Fig. 1C). As expected from the LeX staining (Fig. 1A), both the FUT9-expressing Colo320 and Hutu80 were strongly stained with LTA. As predicted, expression of FUT9 did not affect surface α1-2 fucose (bound by UEA-I) (Fig. 1C). In Colo320, FUT9 ectopic expression did not result in noticeable changes in the overall glycosylation profile (Fig. 1C). In contrast, in Hutu80, FUT9 expression resulted in a small reduction in Tn antigen (*O*-GalNAc, bound by HPA), and α-GalNAc (bound by SBA), as well in α2-6 sialic acid (bound by SNA) in Hutu80-FUT9 compared to Hutu80-Mock, although not statistically significant. These results highlight that glycosyltransferase overexpression under the control of a moderate-strength promoter such as the PGK promoter constitutes a physiologically relevant model for subsequent *in vitro* studies.

### Effects of FUT9 on the expression of MHC-I and CD2 ligands by cancer cells

We next examined whether FUT9 was associated with variation in Major Histocompatibility Complex (MHC) class I surface expression, as low or absent expression of MHC class I is associated with NK cell lysis (Ljunggren & Karre, 1990). We did not observe any differences in MHC class I expression by Mock and FUT9 glycovariants of either Colo320 or Hutu80 (Fig. 2A). We also assessed whether FUT9 would affect the response to Interferon-γ (IFN-γ), a well-known inducer of MHC class I expression (Zhou, 2009). Overnight treatment of cancer cells with IFN-γ increased surface MHC class I compared to steady-state levels (Fig. 2A-B).

**Figure 2.**
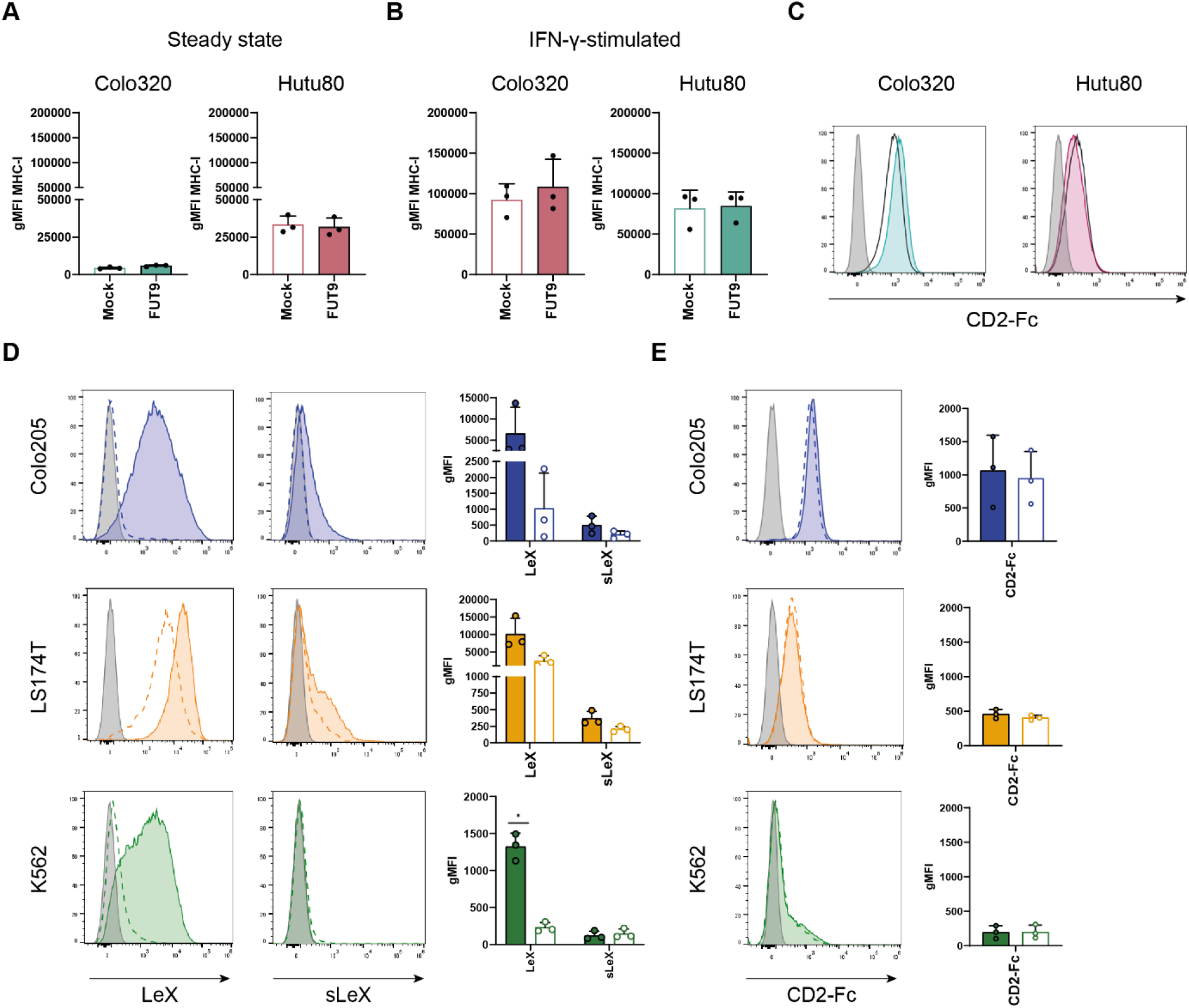
α1-3/4-fucosylation has limited effect on MHC class I expression and CD2 binding. **A-B**, Colo320 (top) or Hutu80 (bottom) glycovariants were stained for surface MHC class I at steady state (A) or after 24 h stimulation with IFN-γ (B) and measured by flow cytometry. **C**, Colo320 (top, green) or Hutu80 (bottom, red) glycovariants were stained with CD2-Fc and analyzed by flow cytometry. One representative experiment. Grey, streptavidin-stained cells; Black, Mock-transfected, Filled, FUT9*-*expressing cells. **D-E**, Colo205 (top, blue), LS174T (middle, yellow), and K562 (bottom, green) were stained for LeX and sLeX (D) or with CD2-Fc (E) after 3 days of treatment with 2F-PF or the DMSO vehicle, and analyzed by flow cytometry. One representative histogram is shown. Grey, secondary antibody; Dashed, 2F-PF-treated, Filled, DMSO-treated. from n = 3; filled bars, DMSO-treated, white bar, 2F-PF-treated. Normality was verified by Shapiro-Wilk and statistical differences calculated with Mann-Whitney U test or unpaired t-test. Not significant unless indicated. *, p < 0.05.

However, surface MHC class I upregulation was not dependent on FUT9 expression in either Colo320 or Hutu80 (Fig. 2B). Altogether, our results show that FUT9 is not involved in regulating MHC class I surface expression, either at steady-state or after IFN-γ stimulation.

LeX has been reported to be a ligand for CD2, a receptor expressed at the surface of lymphocytes, including NK cells (Sabry *et al*., 2011). Both Hutu80 and Colo320 were bound by a recombinant CD2-Fc, consisting of the extracellular domain of human CD2 and the constant fraction (Fc) of human IgG1 (Fig. 2C). Binding of CD2-Fc was independent of FUT9 expression (Fig. 2C), suggesting that LeX epitopes synthesized by FUT9 are not ligands for CD2.

To determine the putative interaction of CD2 with Lewis antigens in a different model, we assessed the binding of CD2-Fc to LS174T and Colo205. Both other colorectal cancer cell lines naturally express LeX and sLeX (Fig. 2D). We also included K562, a lymphoblastic cell line, which was previously used by others to study the interaction of LeX and CD2 (Sabry *et al*., 2011). We found that K562 expressed LeX but sLeX at the cell surface (Fig. 2D). Treatment with the fucosylation inhibitor 2F-PF strongly reduced surface LeX and sLeX in both Colo205 and K562, although this reduction was only significant in K562 (Fig. 2D). Surface LeX could still be detected in LS174T after 2F-PF treatment, albeit at reduced levels (Fig. 2D).

CD2-Fc readily bound the LS174T, Colo205, or K562 cells, yet its binding was unaffected by 2F-PF treatment (Fig 2E), indicating that neither LeX nor sLeX is the ligand of CD2. Of note, 2F-PF also induced a decrease in AAL (binds to both α1-3/4- and α1-6-fucosylated glycans) and UEA-I (binds to α1-2-fucosylated glycans) staining (Fig. S2A), suggesting that 2F-PF also reduced α1-2- and potentially α1-6-fucosylation. Altogether, in contrast to a previously published report (Sabry *et al*., 2011), our data argue against a fucose-based ligand of CD2 and suggest that LeX might actually be dispensable for recognition of cancer cells by NK cells.

### Limited effect of LeX in NK cell-mediated cancer cell destruction and NK cell activation

To determine whether FUT9 and LeX were involved in NK cell-mediated cancer cell lysis, we co-incubated NK cells with cancer cells for either 4h or 24h. However at neither timepoint did we observe any effect of FUT9 on cancer cell killing by the NK cells (Fig. 3A-B).

**Figure 3.**
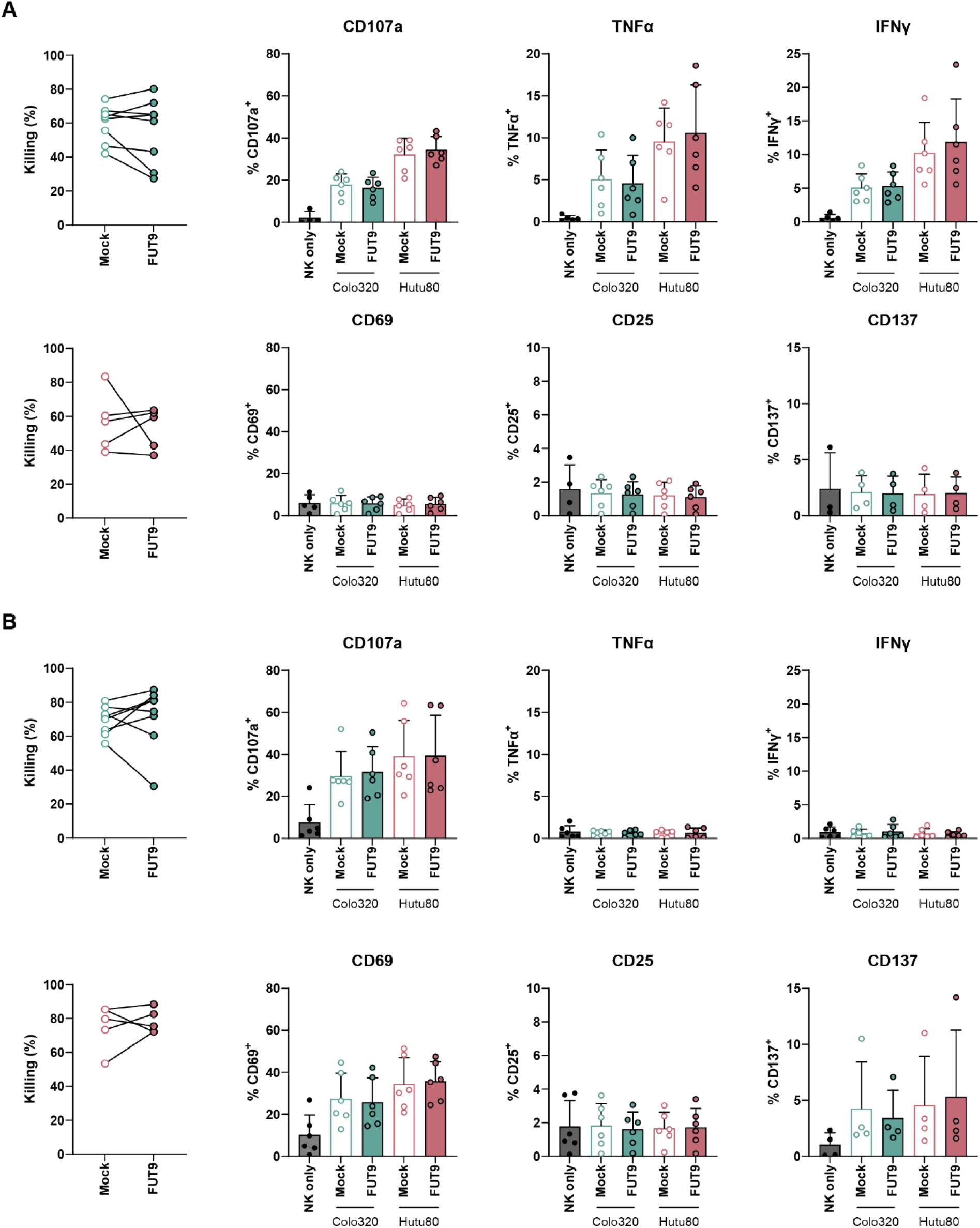
– LeX has a limited role in cancer cells-NK cells interactions. **A-B**, Colo320 (Green) of Hutu80 (Red) glycovariants were incubated with NK cells for 4 h (A) or 24 h (B), after which cancer cell viability was assessed by CTB metabolization. Expression of surface of CD107a, CD69, CD25, and CD137, as well as intracellular TNFα and IFN-γ was assessed by flow cytometry. (n = 5-8 donors, 3-4 independent experiments). Normality was verified by Shapiro-Wilk and statistical differences between Mock and FUT9 conditions were calculated with Mann-Whitney U test or unpaired t-test. All not significant.

We also assessed whether expression of activation markers by NK cells was altered upon co-culture with FUT9 glycovariants. After 4h of co-incubation with cancer cells, we observed an increased proportion of NK cells expressing surface CD107a, a marker of degranulation, as well as intracellular TNFα and IFN-γ (Fig. 3A). However, this increase was independent of FUT9 expression by cancer cells. Similarly, after 24 h of co-culture, we observed an increase in the proportion of CD107a^+^, CD69^+^ and CD137^+^ NK cells, again in a FUT9-independent manner (Fig. 3B). Overall, our data suggest that neither FUT9 nor LeX is a major determinant for NK cell activation and cancer cell elimination by NK cells.

## Discussion

Aberrant glycans produced by cancer cells dampen anticancer immune responses and represent attractive targets for adjuvant therapy to potentiate immunotherapy (Hatinguais *et al*., 2026). Yet, compared to the large body of work characterizing the role of sialic acid-containing glycans in immune evasion (van Vliet & van Kooyk, 2025), our understanding of the role of fucosylated glycans in anticancer immunity remains limited. In this study, we took advantage of the selective activity of FUT9 towards LeX synthesis (Fig. 1A) (Mondal *et al*., 2018) to explore the role of this carbohydrate antigen in the interaction of cancer cells with NK cells.

Using two different colorectal cancer cell lines, we showed that ectopic expression of *FUT9* under the low-to-moderate-strength phosphoglycerate kinase promoter (PGK) (Qin *et al*., 2010) efficiently drove surface LeX expression without major changes in surface glycosylation (Fig. 1A, 1C). FUT9 did not affect the expression of surface MHC class I at steady state nor after IFN-γ stimulation (Fig. 2A-B). In this model, we did not observe any relationship between the presence of LeX and the binding of a soluble CD2-Fc to cancer cells (Fig. 2C, 2E). It has been previously hypothesized that LeX-mediated triggering of CD2 primes NK cells to improve killing of cancer cells (Sabry *et al*., 2011). Interestingly, in their study, Sabry and colleagues found that overexpression of FUT4 led to improved priming of NK cells and that killing assays in the presence of anti-LeX or anti-CD2 antibodies reduced the killing efficiency of NK cells (Sabry *et al*., 2011). Nevertheless, direct evidence of the binding of CD2 to LeX has not yet been established. Our study echoes earlier work showing that the LeX is not a ligand of CD2 (Warren *et al*, 1996), although one specific anti-LeX monoclonal antibody was reported to prevent the binding of CD2 to K562 (Sabry *et al*., 2011; Warren *et al*., 1996). Warren and colleagues reported that CD2 binding to K562 was decreased upon treatment with the bovine pancreatic L-fucosidase (Warren *et al*., 1996). However, treating our cancer cells with 2F-PF, a fucosylation inhibitor that reduced surface α1-2, α1-3/4, and potentially α1-6 fucosylated glycans (Fig. 2D, Fig. S2), did not reduce CD2-Fc binding (Fig. 2E). Nevertheless, we cannot formally rule out that other fucosyltransferases, not evaluated here, could still contribute to synthesizing CD2 ligands on cancer cells.

At a functional level, and in contrast to the effect of sialic acids on NK cell-mediated killing (van Vliet & van Kooyk, 2025), we did not find any role of FUT9/LeX in NK-cell mediated elimination (Fig. 3). These results contradict the earlier work by Sabry et al. (Sabry *et al*., 2011),describing that LeX (and FUT4 expression) were linked to an increased destruction of cancer cells. A limitation in our methodology resides in the overnight incubation of NK cells with IL-2 post-isolation. IL-2 is a known priming agent of NK cells and commonly included in these types of assays (Sabry *et al*, 2019). However, in our hands, we did not observe that the absence of IL-2 priming resulted in a FUT9-dependent effect on cancer cell susceptibility to NK cell-mediated destruction (data not shown).

Altogether, our results suggest that unlike the sialic acid-containing glycans that trigger Siglecs (Hatinguais *et al*., 2026; Hudak *et al*., 2014; van Vliet & van Kooyk, 2025), fucosylated glycans might only have a limited role in the evasion of NK cell-mediated cancer cell destruction. Although glycosyltransferases can also dampen anticancer immune responses by altering the stability of immune evasion-related proteins by altering their glycosylation, we did not find any evidence of such role for FUT9 in our colorectal cancer cell models. Future work will aim at comparing FUT9 with other Lewis antigen-synthetizing fucosyltransferases to evaluate whether other enzymes of this family can promote immune evasion by cancer cells.

## Acknowledgements

The authors thank the Microscopy and Cytometry Core Facility of Amsterdam UMC for their assistance and support.

## Figures and legends

**Figure S1.**
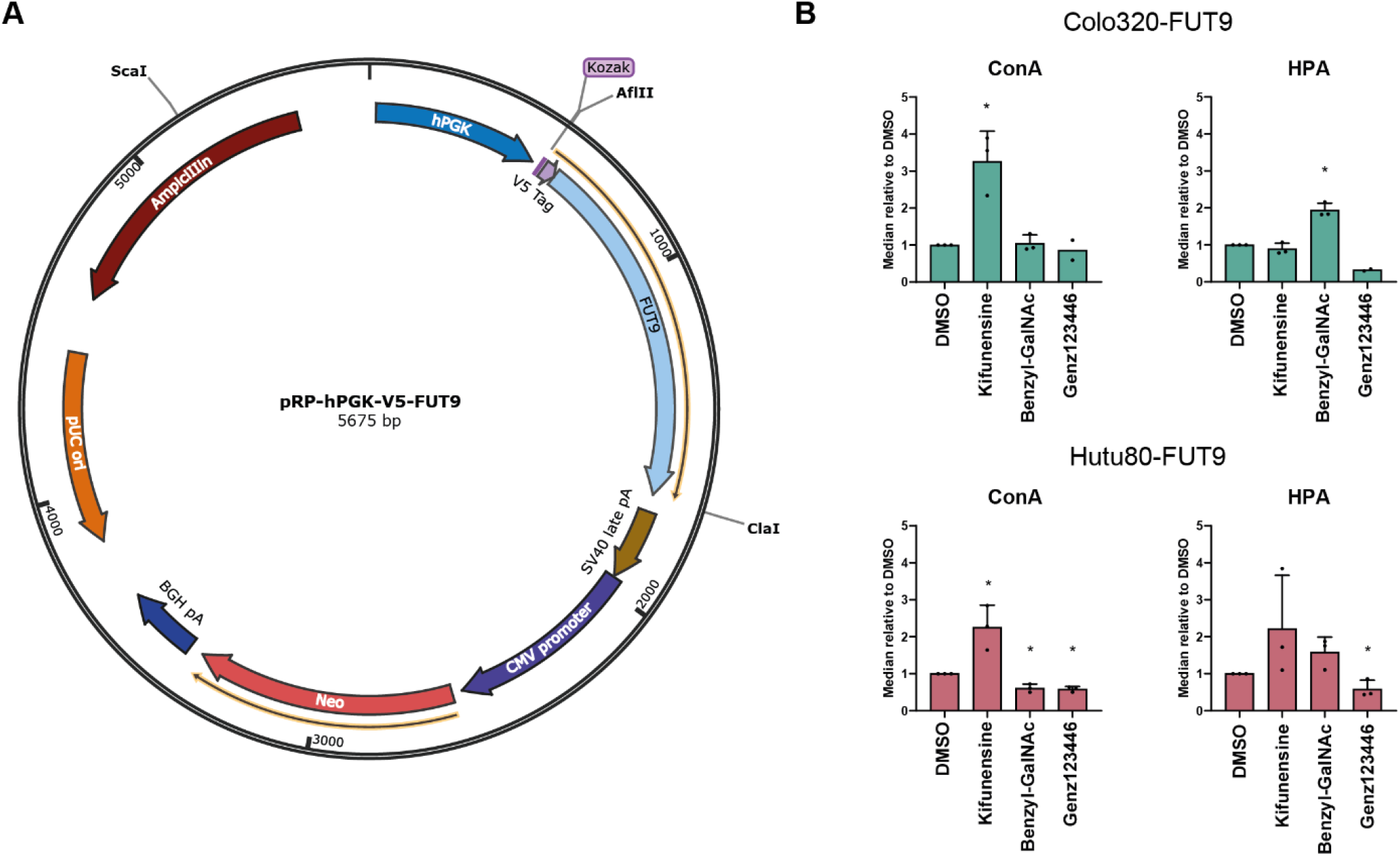
– FUT9-encodind plasmid and effect of glycosylation inhibitors on ConA and HPA staining. **A**, Plasmid map of the FUT9-encoding plasmid. Mock cells were obtained by transfecting the backbone from which *FUT9* was excised using AflII and ClaI. **B**, Colo320-FUT9 (top, green) and Hutu80-FUT9 (bottom, red) were incubated for 72 h with glycosylation inhibitors, and then stained with ConA or HPA and analyzed by flow cytometry. Normality was verified by Shapiro-Wilk and statistical differences calculated with one sample t-test. Statistics were not calculated for Genz123346-treated Colo320-FUT9. *, p < 0.05. n = 2-3.

**Figure S2.**
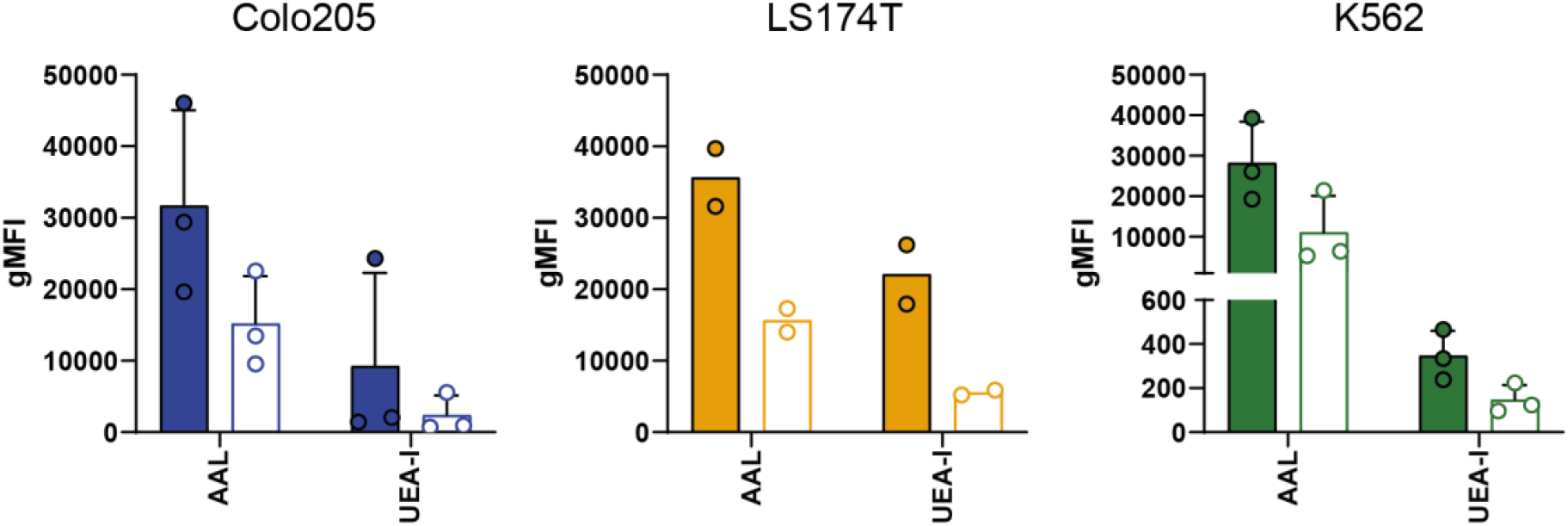
– 2F-PF treatment reduces binding of lectins AAL and UEA-I to cells. Colo205 (left, blue), LS174T (center, yellow), and K562 (right, green) were stained AAL or UEA-I after 3 days of treatment with 2F-PF or DMSO vehicle, and analyzed by flow cytometry. n = 2-3; filled bars, DMSO-treated, white bars, 2F-PF-treated. Normality was verified by Shapiro-Wilk and statistical differences calculated with Mann-Whitney U test or unpaired t-test. All not significant. Statistics were not calculated for LS174T.

